# How does urbanization shape shell phenotype, behaviour and parasite prevalence in the snail *Cornu aspersum*?

**DOI:** 10.1101/2025.01.10.632424

**Authors:** Maxime Dahirel, Youna de Tombeur, Claudia Gérard, Armelle Ansart

## Abstract

Urbanization is a complex and multivariate environmental change, leading to e.g. habitat fragmentation and loss, changes in local climate, soil imperviousness, pollution, etc. This is likely to exert pressures simultaneously on various dimensions of organisms’ multivariate phenotypes, leading to trait shifts with potential ecological consequences. However, responses to urbanization are often studied one (type of) trait at a time. In this context, we studied how, in the brown garden snail *Cornu aspersum*, shell phenotype (shell size and colour/reflectance), behaviour (food intake, mobility) and metazoan parasite prevalence respond to urbanization. Within an urbanization gradient spanning the French city of Rennes, we found that snails from more urbanized sites closer to the urban centre were smaller, whereas urbanization had no detectable effect on shell reflectance, parasite prevalence or behaviour. Larger snails and snails with paler shells were more likely to be infected by trematode metacercariae and sexually-transmitted nematodes (*Nemhelix bakeri*), respectively. Snails harbouring trematode sporocysts ate typically less, while those infected by *N. bakeri* moved more slowly. We discuss the decrease of snail size along the urbanization gradient in relation with the Urban Heat Island effect and the potential decrease of resource quality and availability in urban sites. The absence of detectable effects of urbanization on shell reflectance, mobility and parasite prevalence may be due to scale mismatches between how urbanization is measured and how snails experience microhabitats. We propose further experimental and field studies to decipher interactions between urbanization effects, shell phenotype, life-history traits and parasitism.

## Introduction

Urban areas continue to expand globally (Huang et al. 2019), leading to a wide array of environmental changes including habitat fragmentation and loss, changes in local climate such as the Urban Heat Island (UHI) effect, shifts in water, noise and light regimes, high levels of chemical pollution, etc. (Parris 2016). As a consequence, urban species are usually a non-random sample of regional species pools with respect to traits (Hahs et al. 2023), including body size (Merckx et al. 2018), or mobility and thermal preferences (Piano et al. 2017). The same environmental changes can also lead to phenotypic responses at the within-species level, from plasticity and/or adaptive evolution (Szulkin et al. 2020; Diamond and Martin 2021; Thompson et al. 2022). Urban and non-urban populations of the same species may differ in traits such as body size (e.g. Brans et al. 2017; Martin and Sheridan 2022) or colour (Kerstes et al. 2019; Fukano et al. 2023), but also in more labile traits including movement and other behaviours (Dahirel et al. 2016; Carlen et al. 2021). These within-species phenotypic changes can have wider ecological consequences, as interspecific interactions are often mediated by traits. In non-urban contexts, this has been studied with respect to parasites and intraspecific variation in host traits (Ezenwa et al. 2016; San-Jose and Roulin 2018; Bachtel et al. 2019). Therefore, the effects of urbanization on host-parasite interactions (Murray et al. 2019) might be driven, in part or as a whole, by host phenotypic responses to city life. If this is the case, we should expect host traits and parasite prevalences to simultaneously respond to urbanization, forming an “urban syndrome”.

Terrestrial gastropods (snails and slugs) are good but underused models in urban ecology. Their low active dispersal capacities (Kramarenko 2014) limit their ability to avoid impacts of urbanization by moving, even locally. As ectothermic and humidity-sensitive animals (Cook 2001), urbanization-induced changes in abiotic conditions may influence their behaviour and life history. Compared to other taxa, evidence for within-species trait responses to urbanization is however scarce. Dahirel et al. (2016) showed variation in exploration behaviour between urban and non-urban populations of the snail *Cornu aspersum*. A larger study by Kerstes et al. (2019) used a Netherlands-wide citizen science scheme to show how colour polymorphism in the grove snail *Cepaea nemoralis* responded to urbanization. Shells were generally paler in urban environments, in accordance with previous *Cepaea* studies on the link between colour and climate in other contexts (e.g. Jones 1982; Silvertown et al. 2011; Ożgo and Schilthuizen 2012).

Importantly for studies of species interactions in cities, snails and slugs are hosts to a diverse range of parasites (Barker 2004; O’Brien and Pellett 2022). Infection may affect host traits like feeding or movement behaviour (Dahirel et al. 2022; Rae et al. 2023), which may influence the way snails shape plant communities (Hulme 1996; Frank 2003). The links between parasite infection and behaviour may be correlated with shell phenotype in complex ways. In *Cepaea nemoralis* again, colour morphs differ in baseline behaviour, in infection risk and in their behavioural response to infection (Rosin et al. 2018; Dahirel et al. 2021; Dahirel et al. 2022). In several species, lighter snails are potentially more vulnerable to infection (Scheil et al. 2014; Dahirel et al. 2022), and it has been hypothesized this reflects links between the melanin pathway and immunity, as seen in other invertebrates (Scheil et al. 2013, 2014; Coaglio et al. 2018; San-Jose and Roulin 2018). The mechanism may be different however, as the link between expression of the internal physiological melanin pathway and external shell colour is not clear-cut in snails (Affenzeller et al. 2020).

Furthermore, some parasites infecting terrestrial gastropods are of veterinary and/or human health interest (Giannelli et al. 2016; Butcher 2016; Xie et al. 2024). Understanding how the infection landscape may be reshaped by urbanization, and how this links to host traits, is therefore especially relevant. A synthesis of existing evidence across animal taxa showed that effects of urbanization on parasite infection are highly variable in both magnitude and direction, and taxon-dependent (Murray et al. 2019). While studies of snail parasites or pathogens based on urban samples are not uncommon (Andrus et al. 2022; Xie et al. 2024), they actually rarely investigate the effect of urbanization itself. Nevertheless, a few studies suggest that urbanization can affect parasite dynamics (Aziz et al. 2016; Zhang et al. 2023; Dahirel et al. 2024). However, the direction of effect varied between studies, possibly due to both species effects and differences in the targeted endpoints.

In this context, we focused on the brown garden snail *Cornu aspersum*, a well-studied species common across western European cities. We studied whether body size, shell colour, behaviour (food intake and mobility) and metazoan parasite prevalence simultaneously respond to urbanization in ways consistent with general predictions and previously described trait syndromes. From previous work, we expected more urban snails to be smaller, with lighter shells and potentially more active (Dahirel et al. 2016; Kerstes et al. 2019; Martin and Sheridan 2022). We also expected urbanization to lead to higher parasite prevalence, possibly linked with the lighter shell colour, with negative impacts on behaviour. We explored whether parasite responses may depend on parasite characteristics by investigating several parasite groups and life stages with contrasted infection and transmission strategies.

## Material and methods

### Site selection, snail collection and maintenance

We collected snails between April 12 and April 22, 2022 along an urbanization gradient spanning the northwestern quarter of the city of Rennes (Brittany, western France, ≈ 48°7’ N, 1°40’ W, **Fig. 1**).

**Figure 1.**
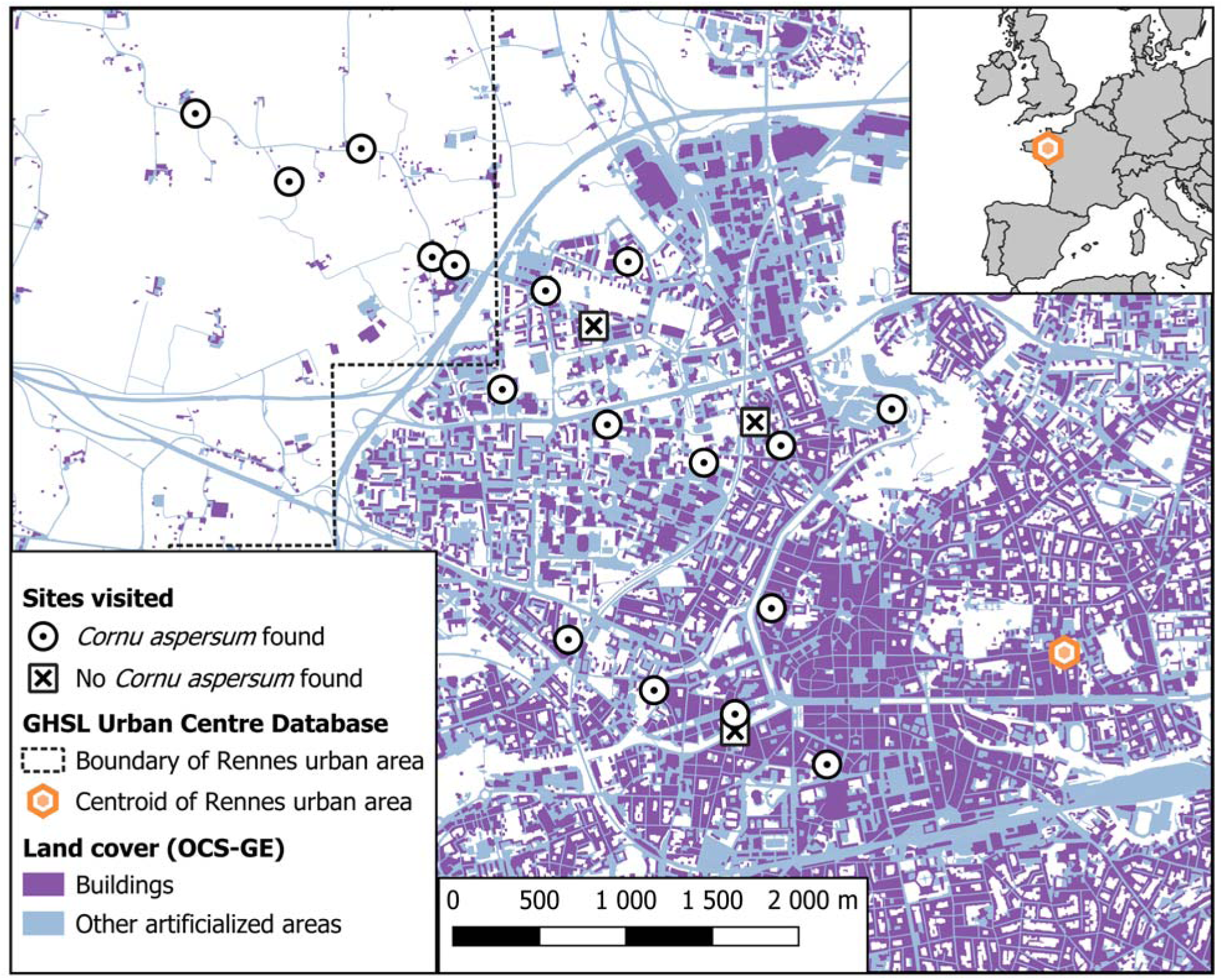
Map of the study area, showing study sites as well as the urbanization gradient, with land cover being less artificialized around sampled sites farther from the urban area centroid. Inset: location of Rennes within France and Western Europe. Map was done using QGIS version 3.38 (QGIS.org 2024). Layers in the main map are from Florczyk et al. (2019) (reused under CC-BY 4.0 license) and Rennes Métropole and Institut national de l’information géographique et forestière (2023) (under Open Database License). Map in inset is from Eurostat (https://ec.europa.eu/eurostat/web/gisco/geodata/administrative-units/countries, licensed under CC-BY 4.0) and is © EuroGeographics for the administrative boundaries.

Although we initially targeted all snails of the Helicidae family, *Cornu aspersum* was the only species found across the entire urbanization gradient (with *Cepaea hortensis* found in only one of the visited sites and *Cepaea nemoralis* in only four, see Gérard et al. 2023), and is therefore the sole focus of the present study. Therefore, while we collected data on all three species (**Data and code availability**), the details, numbers, and analyses described in the rest of the Material and Methods section refer to *C. aspersum* only.

Within that area, we selected publicly accessible sites *a priori* favourable to helicid snails based on personal observations and a database of micro-habitat preferences (Falkner et al. 2001). Twenty sites were prospected, with *C. aspersum* snails found in 17. We chose sites so that populations of *C. aspersum* were separated by at least 100 m, i.e. far enough that they can be safely considered distinct units with respect to gene flow, based on population genetics and known helicid dispersal distances (Arnaud et al. 1999; Kramarenko 2014). In each site, we explored an area of up to 50 m radius around a designated central point, with most of the effort within a 20 m radius. Snails were searched by eye and hand-picked, mainly in tall herbaceous plants used as food sources (in particular *Urtica dioica*, Iglesias and Castillejo 1999) and in or near potential shelters (walls, fences, shrubs…). We only collected adults, easily recognisable by the presence of a reflected “lip” around their shell opening. We visited each site on average 2.55 times (range: 1-5), and collected between 10 and 25 snails per site; for populations where more than 20 individuals of one species were collected across all visits, excess individuals were chosen at random and released back to their capture site. This led us to keep a total of 303 *C. aspersum* individuals from 17 sites.

Snails were housed in plastic boxes with a meshed lid (depending on availability, in either 44.5 × 29.5 × 7.5 cm^3^ boxes with maximum 16 individuals of the same population per box, or in 26.5 × 27.5 × 8.5 cm^3^ with maximum 10 individuals), lined with upholstery foam kept humid, and with additional water and dry snail food in cups *ad libitum* (cereal flour enriched in calcium, Hélinove Performance, La Boupère, France). We gave all snails individualized marks using Posca paint pens (uni Mitsubishi Pencil, Tokyo, Japan), written on the bottom part of the largest shell whorl and covered with transparent nail polish to reduce mark loss. We kept snails under controlled conditions (21 ± 1°C, L:D 14:10 with the dark period aligned with natural night) for at least 10 days between capture and the start of behavioural tests. We cleaned boxes and changed the lining once a week.

### Urbanization metrics

For each site, we used the distance to the centroid of the Rennes urban area (as defined in the GHSL Urban Centre Database version R2019A, Florczyk et al. 2019) as a first measure of the position on the urbanization gradient. To relate this distance to relevant changes in land cover at various spatial scales, we used a nationally standardized and high-resolution land cover layer (OCS-GE, 2020 version for the Rennes area, Rennes Métropole and Institut national de l’information géographique et forestière 2023). Around each site and in buffers of increasing radii from 100, 200, 300, …, up to 1000 m, we collected the proportion covered by buildings (CS 1.1.1.1 in OCS-GE), and by all artificialized surfaces including buildings (CS 1.1 in OCS-GE)(Fig. 1). The initial goal was to determine which land cover metric was the better predictor of urbanization effects, and at which spatial scale. However, we found that for our study sites, built-up cover and total artificial cover were very strongly correlated with themselves across buffer sizes, and each other independently of scale (**Supplementary Material S1**). As a result, the main axis of a PCA including all these land cover variables explained over 93% of the total variation, and was strongly correlated with distance to the city centre (*r* = -0.92)(**Supplementary Material S1**). In practice, this means disentangling the metrics and scales is likely difficult to impossible from our set of sites; therefore, we used distance to the centre of the urban area as our single metric of urbanization for all analyses, with sites closer to the city centre being more artificialized and having a denser built-up cover.

### Behavioural tests

#### Movement speed

All movement tests occurred in the afternoon. Each snail was first placed in a small Petri dish with a few mm of water to stimulate activity; they were then placed individually at the centre of a plastic lid (30 by 30 cm, but with a usable flat area of ≈ 23 by 23 cm). Snails were timed from the moment they left a circle of radius 4 cm drawn around the centre of the lid (this to ensure only times and distances from active snails were counted). In resourceless homogeneous arenas of this size, *C. aspersum* snails tend to move in mostly straight lines after initiating activity (personal observations). We used mucus trails to measure the beeline distance travelled for 90 sec or until they left the plate, whichever happened first; we then used distance travelled and time to calculate the speed of movement away from the starting point. We tested snails twice, with the two tests separated by one week. Six snails died during the experiments, five before movement tests and one between the two movement trials. In addition, three snails never moved during tests, while 15 individuals were only active in one of the trials, resulting in overall 574 speed values from 295 individuals.

#### Food intake

Immediately after each session of movement speed trials, snails were individually placed in small boxes (10 × 10 × 5.5 cm^3^ or 8.8 × 11.3 × 4.1 cm^3^ depending on availability), along with 2 g of dried snail food in a cup. The next day, after 16 hours including the nocturnal activity period, we collected the cups and their content back. We dried the food at 30°C for at least 24 h and weighed it; food intake for each snail was estimated as the difference between the initial 2 g and the remaining flour, recorded to the nearest 0.01 g. We obtained 595 observations from 298 individuals.

### Shell characteristics

After all behavioural tests, snails were killed by freezing at -20°C; after thawing, shells were separated from bodies, and the latter used for parasitological analysis (see below). We used calipers to measure shell size as the shell greater diameter (to the nearest 0.1 mm). Size was measured twice in all snails, non-consecutively. Using the R package *rptR* (Stoffel et al. 2017), we found that size measurements were highly repeatable (*R* = 0.995), and that the estimated residual variation was very small and close to caliper precision (a- = 0.14 mm, or 0.44% of the average size). We therefore used the average of the two shell measurements as a single measure in all subsequent analyses. To measure shell reflectance, we took standardised photographs of cleaned and dried shells in dorsal view (*sensu* Callomon 2019) using a Canon EOS 7D camera. Each photograph included a grey standard card with 9 rectangular cells (7 greys, one white, one black), with the grey scale reflectance values for each rectangle previously determined from spectrophotometry measurements (Maia et al. 2019). We developed an ImageJ/Fiji (Schindelin et al. 2012) and R (R Core Team 2025) pipeline to obtain estimates of shell reflectance as well as their standard errors, using RGB measures of an unworn area of the largest shell whorl and calibration curves based on grey standard cards, following an approach based on Johnsen (2016) (see **Supplementary Material S2** for full details). Here standard errors were not negligible relative to measurements, and were therefore accounted for in statistical analyses (see below and **Supplementary Material S2**).

### Parasitological analysis

As we removed snail bodies, we rinsed shells with water and examined them for metazoan parasites present between the shell and body under binocular microscope (Gérard et al. 2023). We then dissected bodies, and recorded the presence/absence of metazoan parasites infecting the lung, kidney, heart, digestive and reproductive systems. Exact or approximate counts were also recorded for most of these parasites (**Data and code availability**), but were not used in further analyses, as low prevalences and count overdispersion made abundance-based models difficult to estimate. The six snails that were found dead during the experiments (see above) were immediately dissected before decomposition; there are therefore no missing values for parasites. Nematodes found in the reproductive tract were identified as *Nemhelix bakeri* (Morand and Petter 1986; Morand 1988) based on location, adult size and general morphology. *Nemhelix* transmission is very different to all other nematodes known to infect helicid snails, as it is exclusively sexually transmitted (Morand 1988). All other nematodes were pooled together, as they were less frequent (97 nematodes in only 16 snails, excluding a single nematode found in the intestine, where we could not determine whether it was a parasite or an endophoretic nematode, Sudhaus 2018). Land snails can contain trematodes at two larval stages, sporocyst and metacercaria (the latter resulting from infection by cercariae released in the environment by sporocysts) (Butcher and Grove 2001; Gérard et al. 2020). Simultaneous infection by both larval stages can happen, though not by auto-reinfection (Segade et al. 2011; Gérard et al. 2020). We recorded the prevalence of sporocysts and metacercariae separately, especially as their typical target organs (digestive gland and kidney respectively), and therefore potential impacts, are different. Most trematode metacercariae were identified as *Brachylaima* sp., a genus commonly infecting various species of land snails (Segade et al. 2011; Żbikowska et al. 2020; Gérard et al. 2020). A few metacercariae could not be identified conclusively; however, these were always in snails also containing *Brachylaima* metacercariae in greater numbers. We found none of the other metazoan parasites known to infect terrestrial molluscs (Barker 2004).

Shells were also examined for encapsulated parasites, trapped by snails as a defence mechanism (Rae 2017), but only active infections are the focus on the present study. For information on shell-trapped parasites from these individuals, see Gérard et al. (2023).

### Statistical analyses

We analysed our data in a Bayesian workflow (Gelman et al. 2020), using R version 4.5.0 (R Core Team 2025) and the *brms* R package (Bürkner 2018) as frontends for the Stan language (CmdStan version 2.36.0, Gabry et al. 2024; Stan Development Team 2024). For data processing, model post-processing and plotting, we relied mainly on the *tidyverse* (Wickham et al. 2019) suite of packages, along with the *sf* (Pebesma 2018), *bayesplot* (Gabry et al. 2019), *patchwork* (Pedersen 2024), and *tidybayes* (Kay 2023) packages. We used multi-response (generalized) linear mixed models ((G)LMM), which allowed us to estimate simultaneously the effects of our predictors of interest on our response variables, as well as site- and individual-level correlations between the responses (Dingemanse and Dochtermann 2013; Bürkner 2018). We ran three such models: one examining the effects of urbanization on the two measured shell traits, one examining the effects of urbanization and shell traits on the prevalences of the four parasites, and one examining the effects of urbanization, shell traits and parasites on the two behavioural traits.

All continuous predictor variables, as well as response variables in Gaussian models, were centred and scaled to unit 1SD before model fitting to help convergence and prior setting. All models described below make use of the capabilities of *brms* and more generally Stan with respect to measurement error and in-model missing value imputation (Bürkner 2018; McElreath 2020). This allowed us to directly account for measurement uncertainties in reflectance during model fitting, whether it was a predictor or response in a given model.

This also allowed us to fully account for the uncertainty added by missing values in movement speed and food intake. We set weakly informative priors inspired for the most part by McElreath (2020); see **Supplementary Material S3** for full model and prior details. We ran four chains per model, with 3000 warmup and 3000 post-warmup iterations per chain. We checked chain convergence as well as bulk and tail effective sizes were satisfactory following Vehtari et al. (2021). We ran posterior predictive checks to examine model assumptions and confirm the absence of notable mismatch between model and the data (Gabry et al. 2019; Säilynoja et al. 2022). We additionally checked posterior mean model residuals using spline correlograms (Bjornstad 2022) and found no evidence of remaining spatial autocorrelation (95% confidence bands always overlapped with 0). For each model and response variable, we estimated Bayesian R^2^ (Gelman et al. 2019) as overall measures of model fit. We calculated these both excluding and including random effects in predictions; this is analogous, but not identical, to estimating marginal and conditional R^2^ *sensu* Nakagawa and Schielzeth (2013) (the main differences are the way residual variation is computed, and the fact that Bayesian R^2^ are estimated on the observed data scale as opposed to using variance components on the link scale).

For the remainder of this article, all posterior summaries are given as means [95% quantile-based credible intervals].

#### Effect of urbanization on shell phenotypic traits

We analysed shell size and reflectance together using a first multi-response linear mixed model. For both responses, we included distance to city centre as a fixed effect, as well as site identity as a random intercept accounting for other sources of among-site variation. The site-level random effects for the size and reflectance submodels were included in a shared variance-covariance matrix, allowing them to be correlated. Reflectance values are continuous proportions always >0 and < 1, which means a Beta distribution is in principle the most appropriate model (Douma and Weedon 2019). However, we modelled both size and reflectance as Gaussian anyway, which allowed us to estimate residual correlations between the two responses (which reflect here individual-level correlations). This Gaussian approximation is valid here as the observed distribution of reflectance values is far from both the 0 and 1 boundaries, and evaluation of model posteriors confirms that the approximation also doesn’t result in out-of-bounds (::; 0 or :2: 1) predictions (see **Data and code availability** for details).

#### Effect of urbanization and shell phenotype on parasites

We analysed parasite prevalences using a second multi-response GLMM. This model contained four response variables describing active infections: presence/absence of *Nemhelix bakeri*, of other parasitic nematodes, of sporocysts, of metacercariae. All these binary responses were modelled as Bernoulli-distributed with a logit link. For each model, we included urbanization (distance to city centre), shell size and shell reflectance as fixed effects, as well as site identity as a random intercept, again correlated between responses.

#### Effect of urbanization, shell phenotype and parasites on behaviour

We analysed movement speed and food intake together in a third multi-response GLMM. Speed was modelled as a Weibull-distributed variable with a log link (which performed better in posterior predictive checks than lognormal or Gamma models), and food intake as the proportion eaten out of the initial 2 g using a Beta model with a logit link. In both cases, fixed effects included distance to city centre, shell size, shell reflectance, as well as the presence/absence of infections by *N. bakeri*, other nematodes, sporocysts and metacercariae. Random effects included site-specific as well as snail-specific (because of repeated tests) random intercepts, again correlated between the two responses. In 12 cases (out of 595 non-missing values), observed food intake values were 0, which is an issue because Beta distributed variables must always be > 0. To solve this issue, we used one of the transformations suggested by Douma and Weedon (2019): P* = (P(n - 1) + 0.5)/n where P is the original proportion, P* the transformed proportion and n the total number of (non-missing) observations.

## Results

### Effect of urbanization on shell phenotypic traits

Snails coming from more urbanized sites closer to the urban centre were on average smaller, but urbanization had no detectable effect on shell reflectance (**Table 1**, Fig. 2). There was no clear evidence of site-level or individual-level/residual correlation between the two traits (r_[sites]_ = -0.28 [-0.75, 0.29], r_[snails]_ = -0.12 [-0.25, 0.01]). Overall, urbanization and site-level random effects explained only a minority of shell size variation, with most variability in size being within populations; by contrast, variation in shell reflectance was more equally divided into among- and within-site parts (**Table 1**).

**Table 1.** Posterior summary for the model analyzing shell phenotype (with both response variables modeled as Gaussian). See text for values of site- and snail-level correlations between the traits. The full posteriors are displayed in Supplementary Material S4.

|  | Shell size (scaled) | Shell colour (scaled) |
| --- | --- | --- |
| **Fixed effects** |  |  |
| Intercept | -0.01 [-0.20, 0.17] | -0.02 [-0.38, 0.33] |
| Distance to city centroid (scaled) | **0.27 [0.09, 0.46]** | 0.10 [-0.26, 0.46] |
| **Random effects** |  |  |
| Site-level SD | 0.29 [0.11, 0.51] | 0.71 [0.49, 1.05] |
| **Distributional parameters** |  |  |
| Residual SD | 0.94 [0.86, 1.02] | 0.62 [0.56, 0.70] |
| **Bayesian $R^2$** |  |  |
| Full model predictions | 0.14 [0.07, 0.21] | 0.41 [0.34, 0.48] |
| Fixed effects only | 0.08 [0.01, 0.18] | 0.04 [0.00, 0.16] |

**Figure 2.**
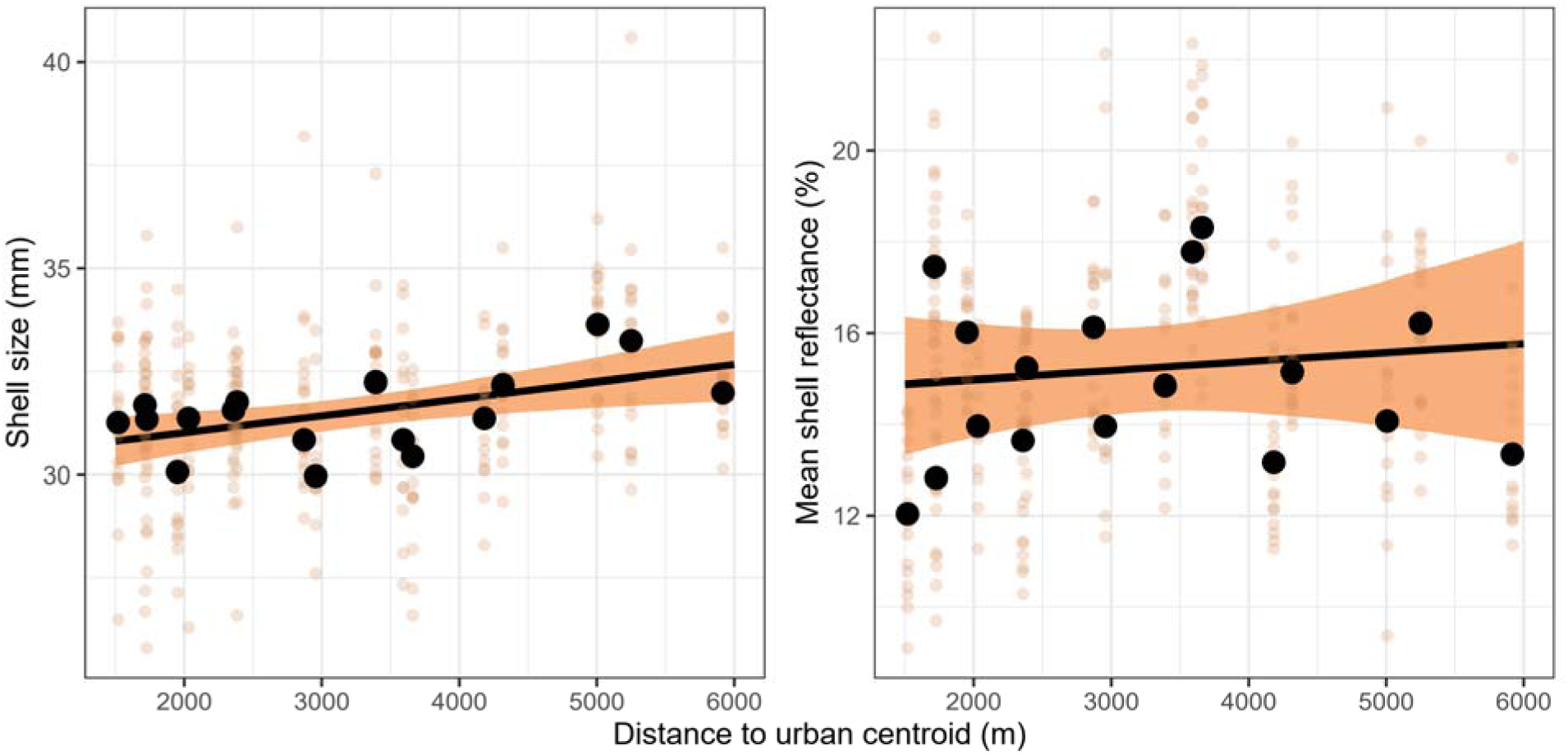
Effect of urbanization (measured as distance to city centroid) on shell phenotype (shell size and colour). Transparent points are individual observed values, large black dots are observed population means, while black lines and coloured bands are posterior mean model predictions and their 95% credible intervals (based on fixed effects). Measurement uncertainty in individual shell reflectance values is not displayed, to keep the plot readable, but is accounted for in the model; predictions are made assuming the average reflectance uncertainty.

### Effect of urbanization and shell phenotype on parasites

Urbanization had no detectable effect on the prevalences of any of the investigated parasite groups (*Table 2*). Larger snails were more likely to be infected by metacercariae, while snails with paler shells (higher reflectance) were more likely to harbour *N. bakeri* infections (*Table 2*, Fig. 3). There was no clear support for site-level correlations between any of the parasites (*Table 3*). Based on Bayesian R^2^ values, the proportion of variation in parasite prevalence explained by shell phenotype and urbanization was limited, even for metacercariae and *N. bakeri* (*Table 2*). However, R^2^ including site random effects were substantially higher, indicating between-sites variation in prevalence not linked to the urbanization metric (*Table 2*; see also *Supplementary Material S5*, for a breakdown of observed prevalences per parasite and population).

**Table 2.**
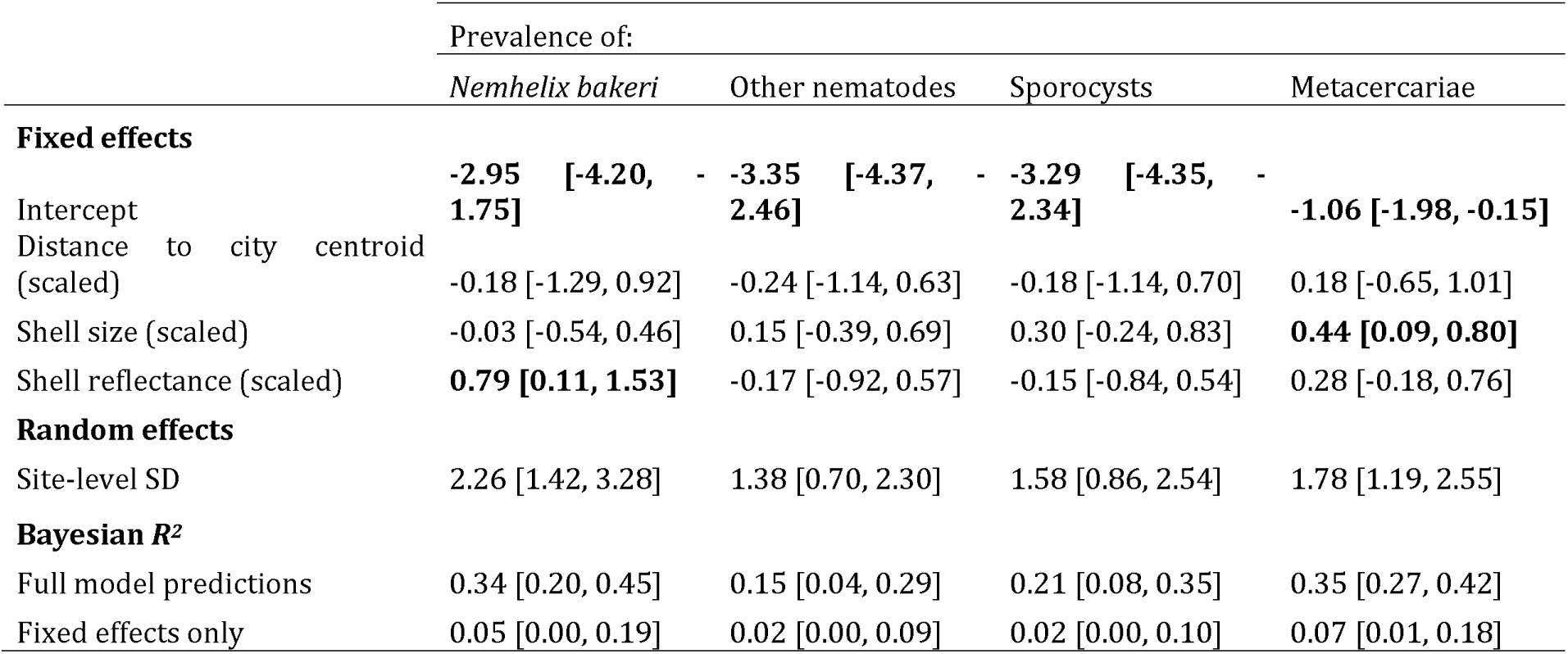
Posterior summary for the model analyzing parasite prevalence (all responses modeled as Bernoulli-distributed). The full posteriors are displayed in Supplementary Material S4.

**Figure 3.**
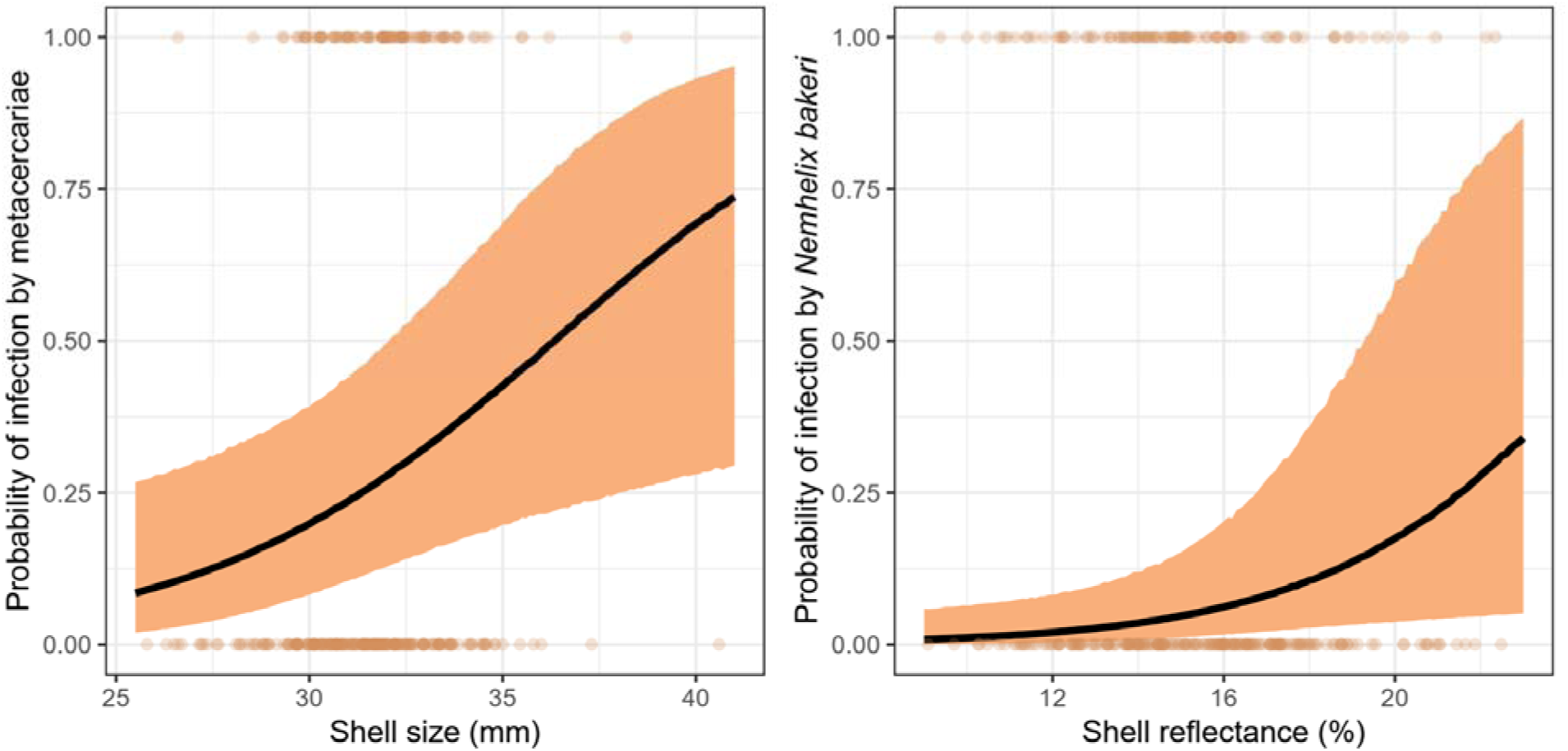
Left: effect of shell size on the probability a snail harbours trematode metacercariae; right: effect of shell colour (average reflectance) on the probability a snail is infected by the nematode Nemhelix bakeri. Transparent points are individual observed values, while black lines and coloured bands are posterior mean model predictions and their 95% credible intervals (based on fixed effects). Measurement uncertainty in individual shell reflectance values is not displayed, to keep the plot readable, but is accounted for in the model; predictions are made assuming the average reflectance uncertainty, and assuming average values for predictor variables not in the plot.

**Table 3.** Posterior summary of the site-level correlations between parasites. The full posteriors are displayed in Supplementary Material S4.

|  | *Nemhelix bakeri* | Other nematodes | Sporocysts |
| --- | --- | --- | --- |
| Other nematodes | 0.25 [-0.26, 0.70] |  |  |
| Sporocysts | 0.46 [-0.00, 0.83] | 0.09 [-0.47, 0.61] |  |
| Metacercariae | 0.10 [-0.34, 0.51] | 0.30 [-0.22, 0.74] | -0.11 [-0.57, 0.38] |

### Effect of urbanization, shell phenotype and parasites on behaviour

Urbanization and shell reflectance had no detectable effect on either movement speed or food intake (*Table 4*). Larger snails typically ate more (*Table 4*, Fig. 4) but were not faster or slower (*Table 4*). Some infections were linked to behavioural differences: snails infected by *N. bakeri* were on average slower (*Table 4*, Fig. 5), while snails infected by sporocysts in the digestive gland ate typically less (*Table 4*, Fig. 5). Food intake and movement speed were not clearly correlated at the site level (r_[sites]_ = 0.52 [-0.00, 0.87]) or the individual level (r_[snails]_ = 0.20 [-0.59, 0.83]). Based on R^2^ values, the model explained only a limited part of movement speed variation, which given model structure indicates that most movement variation is at the within-individual level; by comparison the model explains over half of observed variation in food intake (*Table 4*). The proportion of the variation explained by the model that is attributable to fixed effects specifically is similar (roughly a third) between movement speed and food intake (*Table 4*).

**Table 4.** Posterior summary for the model analyzing snail behaviour (movement speed is modeled assuming a Weibull distribution, food intake assuming a Beta distribution). See text for values of site- and snail-level correlations between the traits. The full posteriors are displayed in Supplementary Material S4.

|  | Movement speed | Food intake |
| --- | --- | --- |
| **Fixed effects** |  |  |
| Intercept | -0.03 [-0.15, 0.08] | **-1.11 [-1.34, -0.87]** |
| Distance to city centroid (scaled) | 0.02 [-0.09, 0.13] | -0.13 [-0.35, 0.10] |
| Shell size (scaled) | -0.02 [-0.07, 0.04] | **0.22 [0.15, 0.30]** |
| Shell reflectance (scaled) | 0.03 [-0.04, 0.10] | -0.02 [-0.13, 0.08] |
| *Nemhelix* present | **-0.33 [-0.52, -0.14]** | -0.07 [-0.34, 0.21] |
| Other nematodes present | -0.09 [-0.30, 0.14] | -0.25 [-0.60, 0.08] |
| Sporocysts present | -0.07 [-0.29, 0.15] | **-0.55 [-0.88, -0.23]** |
| Metacercariae present | -0.03 [-0.15, 0.08] | -0.06 [-0.23, 0.10] |
| **Random effects** |  |  |
| Site-level SD | 0.19 [0.12, 0.30] | 0.44 [0.29, 0.65] |
| Snail-level SD | 0.07 [0.00, 0.18] | 0.39 [0.31, 0.46] |
| **Distributional parameters** |  |  |
| Shape parameter | 1.96 [1.82, 2.11] | -- |
| Precision parameter | -- | 16.08 [13.70, 18.69] |
| **Bayesian $R^2$** |  |  |
| Full model predictions | 0.15 [0.09, 0.24] | 0.59 [0.53, 0.63] |
| Fixed effects only | 0.05 [0.02, 0.10] | 0.16 [0.09, 0.26] |

**Figure 4.**
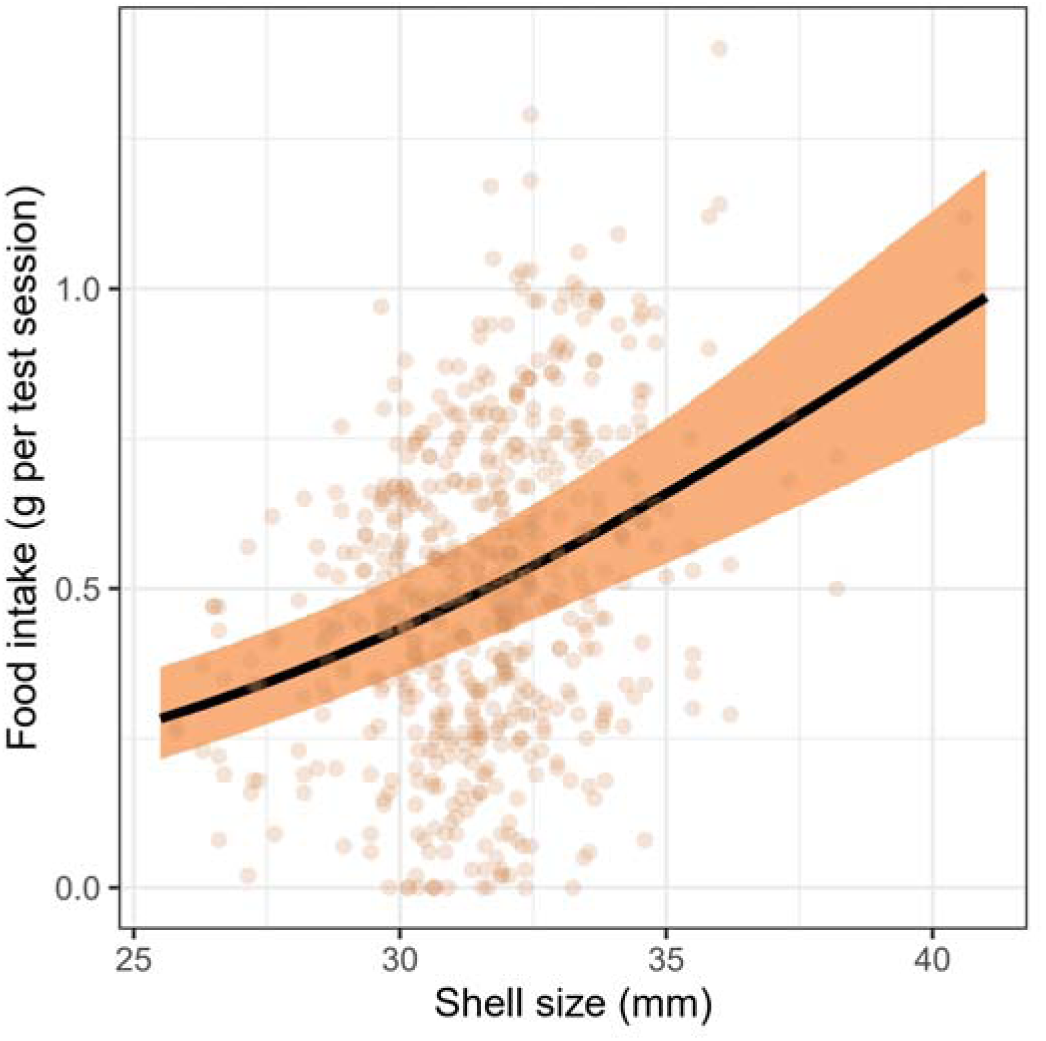
Effect of shell size on snail food intake. Transparent points are observed values for individual trials, while black lines and coloured band are posterior mean model predictions and their 95% credible intervals (based on fixed effects). Predictions are made assuming uninfected individuals as well as average shell reflectance and urbanization; predicted food intake is back-transformed from proportions to grams for plotting.

**Figure 5.**
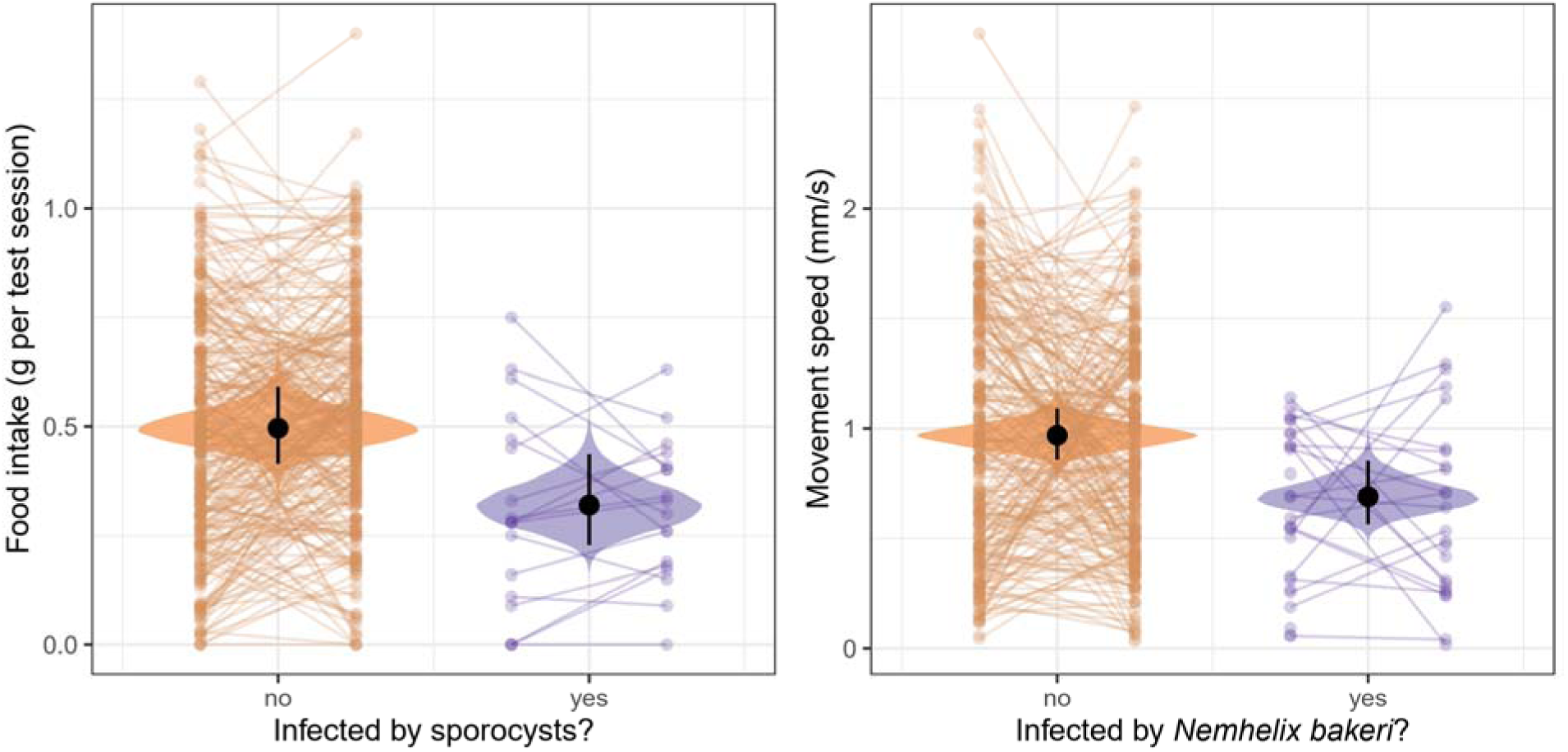
Left: effect of trematode sporocyst infection on food intake; right: effect of Nemhelix bakeri infection on movement speed. Transparent points are observed values for individual trials, with consecutive trials from the same individual connected by lines. Posterior predictions are displayed as violins, with large dots and black bars being posterior means and their 95% credible intervals (based on fixed effects). Predictions are made assuming individuals are uninfected except for the parasite relevant to the plot, and assuming average shell size, shell reflectance and urbanization; predicted food intake is back-transformed from proportions to grams for plotting.

## Discussion

### Cornu aspersum body size decreases along the urbanization gradient

The only clear effect of urbanization we were able to detect was that *Cornu aspersum* adult snails were smaller closer to the city centre. Our results here are consistent with previous anecdotal observations in the same city and species (Dahirel 2014). Shifts toward smaller sizes in cities have been reported at both community and within-species level in several taxa (e.g. Brans et al. 2017; Merckx et al. 2018; Dahirel et al. 2019; Braschler et al. 2021; Martin and Sheridan 2022). They are often attributed to the Urban Heat Island effect through Atkinson’s temperature-size rule (Atkinson 1994; Martin and Sheridan 2022), referring to a general trend in ectothermic species to a decrease in size with increasing temperature. This may be mediated both through developmental plasticity (through the temperature dependency of metabolism and growth rate) or through selection, as individuals differing in size may have different thermal tolerances (due to e.g. surface/volume effects with respect to heating or cooling) (Atkinson 1994). While we did not measure temperature directly in this study, the UHI effect over the city of Rennes is well characterized (Foissard et al. 2019) and depending on season, can reach up to 5°C on average. In the helicid snail *Theba pisana*, heating and cooling rates are strongly size-dependent, suggesting that climate may exert selective pressures on body size (Knigge et al. 2017). However, in *C. aspersum* specifically, the evidence for the direct applicability of the temperature-size rule is more mixed. In experimental settings, *C. aspersum* snails reared under warmer treatments reached larger, not smaller, adult sizes than those reared in colder contexts (Czarnoleski et al. 2016). By contrast, there was no clear difference in shell size between populations spread across a latitudinal gradient in Chile, but higher latitude snails were nonetheless heavier (Gaitán-Espitia et al. 2013).

The other commonly advanced explanation for smaller sizes in cities is the diminished quality and/or availability of food (Martin and Sheridan 2022). The increased fragmentation of green spaces in highly urbanized habitats might reduce feeding choices; higher intrapopulation competition for limited quality resources in smaller urban fragments could also lead to reduced size; growth restriction due to intra- and inter-specific competition for resources and/or space is well-documented in helicid snails (e.g. Cameron and Carter 1979; Dan and Bailey 1982). Smaller sizes under sustained food limitation may result from both plastic and evolved responses, as seen in e.g. *Drosophila* (Kolss et al. 2009). While overall a generalist species, *C. aspersum* shows very strong feeding preferences for stinging nettles *Urtica dioica*, a plant with high protein, ash and calcium content, which also represents a suitable habitat (Iglesias and Castillejo 1999). The relationship between snails and nettles could be altered in urban environments for instance due to pollution; *Cepaea nemoralis* exposed to heavy metal polluted nettles reduced their consumption (Notten et al. 2006). Experiments manipulating food access of snails from both urban and non-urban sites during development may help confirm the role of resource quality and availability on our observed results. However, *C. aspersum* time to maturity can reach over a year (Ansart et al. 2009), especially in conditions of resource competition (e.g. Dan 1978), limiting the practicality of such experiments. On the other side of the trophic interactions, snails are often predated in a size-dependent way by rodents and birds (Rosin et al. 2011), with the direction of selection (preferring smaller or larger individuals) depending on both the snail and predator species. Changes in predator communities along urbanization gradients could therefore lead to changes in snail sizes. Correlative approaches associating wild snail measurements to assessments of both the surrounding vegetation and its palatability (Dan 1978), as well as predator communities, may here prove helpful to understand the role of trophic interactions in shaping the size of urban snails.

### Shell reflectance and snail movement speed are not influenced by urbanization

Against expectations, we found no link between shell reflectance variation and the urbanization gradient. Drivers and consequences of genetically-based shell colour polymorphism in gastropods have been the subject of many ecological and evolutionary studies (Jones et al. 1977; Rosin et al. 2013; Williams 2017; Cook 2017; Schweizer et al. 2019; Saenko and Schilthuizen 2021; Gefaell et al. 2021). In several land snail species, pale morphs are often more common in generally warmer than cooler conditions (Schilthuizen 2013; Cook 2017; Köhler et al. 2021). In *C. nemoralis*, in which colour variation is usually seen as clear discrete morphs under strong genetic determinism (but see Davison et al. 2019), a previous study showed that lighter morphs were generally more frequent in urban populations, and explicitly linked this to the UHI effect (Kerstes et al. 2019). In *C. aspersum*, shell colour variation is effectively continuous, resulting from the combination of variation in background colour, in the size, number and colour of spiral bands superimposed over it and on whether and how these bands are mottled or speckled (Chevallier 1977). The overall reflectance we measured is therefore under a complex mix of genetic influences (Albuquerque de Matos 1984b,a), but is also subject to environmental determination. In experimentally bred *C. aspersum*, Lecompte et al. (1998) showed that snails reared for three months at 25°C had significantly lighter shells than those reared at 15°C, as expected from predictions on the link between colour and temperatures (see above). However, the authors did not follow snails up to maturity and warned that these differences could be largely attenuated in adult snails. Further studies examining the colour of immature urban and non-urban snails, either from wild populations or from experimental animals reared under controlled climatic conditions, may be important to clarify whether and how urban climate influence colour variation. Such studies may include more detailed investigation on not just average shell darkness but also colour patterns, especially as these may also interact with thermal tolerance in complex ways (Knigge et al. 2017; Kerstes et al. 2019).

While the urbanization gradient did not explain variation in shell colour, there were nonetheless consistent inter-sites differences in shell reflectance. Climate variables are not the only environmental parameters influencing shell colour variation (Cook 2017), and within-city variation in e.g. predation risk might contribute to the observed results. Furthermore, while our urbanization metric describes habitat at a relatively broad scale (see buffer widths in *Material and Methods*), snails actually experience their surroundings over much smaller areas (with home ranges of a few m^2^, Dan 1978). Between-population differences may be even larger than we quantified as our sampling protocol, which involved search radius closer to maximal dispersal distances than snail home ranges (Kramarenko 2014), may have mixed snails from different micro-habitats together. In order to get a fuller picture, future studies examining *C. aspersum* colour variation in cities may need to both sample individuals and measure their environment at the spatial scale actually experienced by snails.

In a previous study we also led in the city of Rennes (Dahirel 2014; Dahirel et al. 2016), we found that exploration propensity of adult *C. aspersum* increased in urban population to the levels of subadults (the most dispersive stage), suggesting an advantage in maintaining this costly trait in fragmented urban areas. Although this is not immediately contradictory, in the present study adult movement speed did not vary with urbanization. Our movement speed measure, assessed on a very short experimental time, may not be a good proxy for dispersal ability on larger time and spatial scales. In addition, we here measured speed and used a uniform substrate with no variation over the test area and no variation between populations; by contrast, Dahirel et al. (2016) measured boundary-crossing propensity in semi-natural conditions at the boundary between heterogeneous natural and artificial substrates. Pembury Smith and Ruxton (2021) examined movement speed in *C. aspersum* using a similar protocol to ours, with few key differences. While they, like us, did not find any relationship between body size and speed, they showed that crawling speed in *C. aspersum* depended on the substrate texture and orientation, with abrasive substrates significantly slowing down individual snails. Understanding differences in movement between urban and non-urban snails may require investigating their behaviour across a range of substrates with different textures and absorption properties (McKee et al. 2013), covering the diversity of substrates snails may encounter in cities. It may also require studying changes in movement properties beyond speed; organisms may alter site fidelity or path tortuosity in response to habitat fragmentation or matrix harshness (Cote et al. 2017). More generally and as with shell colour, movement differences between populations may be more dependent on local microclimatic and microhabitat parameters as than on the broad scale urbanization gradient (Rosin et al. 2018) (even though the latter will influence the former).

### Parasites are not influenced by urbanization, but snail phenotypic responses and infection are linked to each other

We did not find any clear trend in the prevalence of any studied parasite group along the urbanization gradient, despite substantial inter-site differences in prevalence. This result was especially unexpected for non-sexually transmitted nematodes, as Dahirel et al. (2024) found that *C. nemoralis* snails were less likely to show encapsulated nematodes in their shell (a defence mechanism) in Belgian urban sites. They suggested that among other mechanisms, this may stem from reduced infections in urban populations, themselves linked to lower abundance of soil nematodes in urban areas (Gong et al. 2024; Hu et al. 2024), as many parasitic nematodes known to infect land snails have at least one free-living stage in the soil (Grewal et al. 2003; Morand et al. 2004). In contrast with these findings, Aziz *et al*. (2016) found that in the UK, urban slugs were more, not less, infected by the nematode *Angiostrongylus vasorum*. The link, if and when it exists, between urbanization and non-sexually transmitted nematode infection is therefore more complex than expected, and may involve variation in snail immunity and various epidemiological factors (e.g. occurrence of other invertebrate and vertebrate hosts) (Dahirel et al. 2024).

The absence of a detected effect urbanization on parasites with no lasting free-living stage, such as the sexually-transmitted nematode *N. bakeri* or trematodes, is less surprising, as these parasites may be more primarily constrained by their host presence and characteristics than environmental conditions. For *N. bakeri* in particular, high inter-site variability may reflect the history of direct transmission of parasites between sexual partners in slow-moving host species (especially given the possible link between movement and infection, see below). Although to a lesser extent, the site effect on the prevalence of trematodes (sporocysts and metacercariae) could be related to the abundance of their definitive vertebrate hosts. In a study in Ukraine, contamination of soil with helminth eggs in urban park areas was very heterogeneous and on average twice higher than in rural counterparts, possibly in part due to infected domestic dogs and cats (Paliy et al. 2019). Indeed, cities are often heterogeneous (Alberti et al. 2020) and this spatial heterogeneity, sometimes driven by within-city social dynamics, might end up being a major underappreciated driver of species interaction patterns (Martin et al. 2024).

At the individual level, some parasite prevalences were nevertheless dependent on snail shell traits. In particular, we found that trematode metacercariae were more likely to be found in larger snails. *Brachylaima* metacercariae are pathogenic for *C. aspersum* because they directly feed on the renal epithelium (Segade et al. 2011). Our result could be explained by a better survival of larger snails to these pathological effects as demonstrated for the freshwater gastropod *Bithynia tentaculata* (Bachtel et al. 2019), and/or by a higher infection rate of large snails, offering more biomass, energy and space to the parasites (Poulin and George-Nascimento 2007). Further studies including experimental infections (Segade et al. 2011) may here provide answers. The sexually transmitted nematode *N. bakeri* had a higher prevalence in paler snails, although shell colour had no influence on other types of parasites. There is a bundle of evidence linking dark phenotypes to a better defence system in some land snail species: dark and/or banded morphs are less parasitized (Cabaret 1983, 1988; Scheil et al. 2014; Dahirel et al. 2022) and more prone to wound healing (Scheil et al. 2013). The melanin pathway, through phenoloxidase activity, plays a role in snail immune defence (Coaglio et al. 2018). However, the mechanisms underlying the link between dark phenotypes in snails and defence remain unclear, especially since Affenzeller et al. (2020) showed that dark bands in the polymorphic *C. nemoralis* were not related to melanin production. Both *N. bakeri* prevalence and shell reflectance show high between-population variability; we re-examined these data and found no clear *between*- population correlation between the two (r_[sites]_ = -0.30 [-0.70, 0.18]; *Supplementary Material S6*). While there are obvious limitations due to the low number of populations with non-zero *N. bakeri* prevalence, this suggests that the effect of reflectance on *N. bakeri* seen in the main model could be the result of within-population variability rather than between-population differences. Here again, experimental infections could help understand the link between infection and shell colour, although such protocols are much more difficult to put in place and invasive (Morand and Faliex 1994), compared to trematode infections.

While regarding the results above, it is easy to assume that shell phenotype influences adult infection, rather than the inverse, the direction of the causal links between infection and behaviour may be more difficult to ascertain. Indeed, while consistent individual differences in host behaviour may influence infection risk, infection may itself alter host behaviour, leading to complex feedbacks (Ezenwa et al. 2016). Different parasite infections were associated with behavioural variation:

- Snails infected with sporocysts ate less on average. The effect of parasites on food consumption has been studied in many host-parasite pairs, with results going in either direction; however, parasites directly infecting digestive organs are expected to reduce feeding rates, including through direct physical impediment to the feeding and digestive functions (Mrugała et al. 2023). While *Brachylaima* metacercariae typically infect snails’ kidneys, sporocysts develop in the connective tissue of the digestive gland, which can be totally invaded and destroyed by the sporocyst network (Segade et al. 2011; Gérard et al. 2020). Based on this, our results are here fully consistent with Mrugała et al. (2023)’s synthesis.
- Snails infected with *N. bakeri* were also less active. Parasites often have negative impacts on host movement (Binning et al. 2017) and *N. bakeri* has strong negative effects on fitness (Morand 1989). However, as this nematode is sexually transmitted, older snails, with longer mating histories, may be more likely to be infected. While a direct relationship with age was not proved, there exists some evidence that subadults/young adult snails are more mobile than older adults, including in *C. aspersum* (Tomiyama and Nakane 1993; Dahirel et al. 2017). The question remains therefore open whether *N. bakeri* truly reduces snail activity, or whether the observed association between the two is a by-product of snail age.

### Conclusion and perspectives

Our study showed that urbanization influences some aspects of *Cornu aspersum* phenotype and that phenotype and parasite infection are linked, but was not able to link phenotypic traits, parasite prevalence and behaviour in an “urban syndrome”. There were however substantial between-sites differences (in particular for shell traits and parasite prevalence) despite high intrapopulation variation. This importance of intrapopulation variation may reflect the need to consider environmental variation more adequately at the scale experienced by snails (Bergey 2019). Between-population differentiation in traits could also be the outcome of local population genetic divergence and genetic drift caused by urbanization and fragmentation at local scale (Balbi et al. 2018). While urban population genetic investigations are common (Diamond and Martin 2021), studies simultaneously examining phenotypic divergence and genetic divergence between populations remain rare and may prove fruitful (Rivkin et al. 2019). Finally, it is important to remember that several of the phenotype-parasite associations we describe here are based on low prevalences (*Supplementary Material S5*), and that further studies are likely needed to understand the effects of parasite infections on (urban) snails. New methods of investigation, including soil eDNA (Jaffuel et al. 2019), or examining shells for encapsulated parasites (Rae 2017; Gérard et al. 2023; Dahirel et al. 2024) may be here very useful. Nonetheless, the interdependence of parasite prevalences, food intake and body size, when the latter is itself dependent on urbanization, may open paths for complex indirect links between urbanization and infection dynamics even in the absence of direct effects.

## Acknowledgments

We thank Bram Vanthournout, who provided us with the grey standard cards along with their reflectance values, and Matthew Shawkey, who provided access to the spectrometer used for the calibration of these cards.

## Author Contributions

Initial idea: MD, CG, AA; site selection and fieldwork: MD, YdT, CG, AA; behavioural tests, parasitological analysis, shell size measurements and shell photographs: YdT, CG, AA; data analysis: MD, with preliminary analyses by YdT; initial manuscript draft: MD, based on a preliminary report by YdT.

## Ethics statement

This study complied with all relevant international and national laws, and followed the ASAB/ABS Guidelines for the use of animals (ASAB Ethical Committee and ABS Animal Care Committee 2022) as closely as possible. No ethical board recommendation or administrative authorization was needed to work with, or sample, the study species. The marking method used is non-invasive and has minimal to no documented effects on life-history traits (Henry and Jarne 2007).

## Data and code availability

Data and code needed to reproduce all analyses presented in this article are available on GitHub (https://github.com/mdahirel/urban-snail-parasites-2022) and archived in Zenodo (DOI: https://doi.org/10.5281/zenodo.14628820).

