## Supplementary Material for "How does urbanization shape shell phenotype, behaviour and parasite prevalence in the snail *Cornu aspersum*?"

### S1 - Correlations between urbanization metrics

In the main text, we described how we used distance to the city centroid as our only urbanization variable, as it was strongly correlated with built-up/artificial surfaces % at all scales of interest and these were in turn strongly correlated with each other. We explore this in more detail here.

Correlation coefficients $r$ between any two built-up/artificial surfaces metrics (at any buffer width considered) ranged from 0.73 to 1 with an average $r$ of 0.93. When focusing only on correlations between built-up values at different buffer widths, the distribution of correlation coefficients is similar (0.75 to 1, with a mean of 0.94). When considering only artificial surfaces % values, correlations are even higher (0.88 to 1, with a mean of 0.97).

Considering this, approaches attempting to find a best land cover metric and a best scale to use in our analyses were likely to fail, as metrics and scales would be likely difficult to disentangle from each other. We ran a Principal Component Analysis and found indeed that the first axis of the PCA explained most (93.17%) of the variation in our land cover data, and that all metrics were strongly correlated to it (**Fig. S1-1**; mean correlation of included metrics with PC1: 0.96, range: 0.83 to 0.99).

Additionally, while it was **not** included in the PCA, distance to the urban centroid was nonetheless also strongly correlated to that first PC axis ($r$ = -0.92). We therefore used distance to the city centroid as the sole urbanization variable in our analyses.

**Figure S1-1.** Principal Component Analysis of urbanization-related land cover variables (built-up % and all artificialized surfaces %) for the 17 sites (out of 20 prospected) where *Cornu aspersum* snails were found.

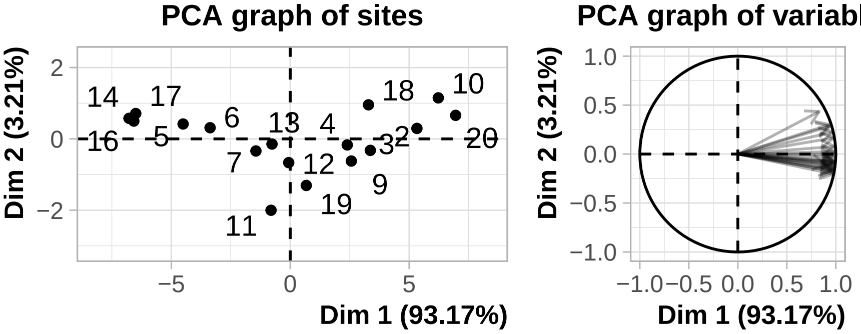


Figure 1

### S2 - Reflectance measurements - extended methods

As mentioned in the main text, we measured shell reflectance using standardised photographs of cleaned and dried shells placed near a grey standard card with 9 rectangular cells (7 greys, one white, one black, **Fig. S2-1**), with the grey scale reflectance values for each rectangle previously determined using spectrophotometry.

To obtain these reference values, the diffuse reflectance was measured using a AvaSpec-2048 spectrometer and a dual light source set-up (AvaLight-DH-S deuterium halogen and AvaLight-HAL-S-MINI light source) with a bifurcated probe and integrating sphere (AvaSphere-50-REFL). A white reflectance standard (WS-2, Avantes) and black standard (BS-2, Avantes) were used for standardization. For five randomly selected cards, three replicates of each rectangle per card were measured. Reflectance spectra were processed using the *pavo* R package (Maia et al. 2019), and mean reflectance obtained for each grey cell by averaging the values for the three replicates of each card, then averaging these values across the five replicate cards (see **Data and code availability** in the main text for resulting values).

We converted white balanced RAW images to TIFF files to use in ImageJ/Fiji (Schindelin et al. 2012), and for each image, we measured the average RGB values of each cell of the grey standard, along with an area of interest on the snail shell (see below). We used values from the grey standards to fit exponential calibration curves for each channel of each individual image, linking the average RGB values of an area of interest to its average grey scale reflectance (Johnsen 2016). All R, G and B calibration curves had very good performances (mean correlation between observed and predicted: 0.998; range: 0.986 - 1).

Periostracum, the outer pigmented layer of the shell, typically gets increasingly damaged and worn as snails age (Williamson 1979). Because of this, we did not measure the average RGB values over the entire shell, as they would not be an accurate representation of colour as it was produced by the snails. We instead measured mean RGB values of an unworn rectangular area covering the entire height of the largest (= most recent) whorl (**Fig. S2-1**). The selected rectangle was parallel to growth lines (which are at an angle compared to the columellar axis) and as close as possible to the columellar axis; its exact positioning and width varied so that no areas with worn periostracum were included. This had the added benefit of avoiding areas with light reflections and the presence of the individual paint marks in some images. We used the average predicted reflectance across RGB channels in our further analyses.

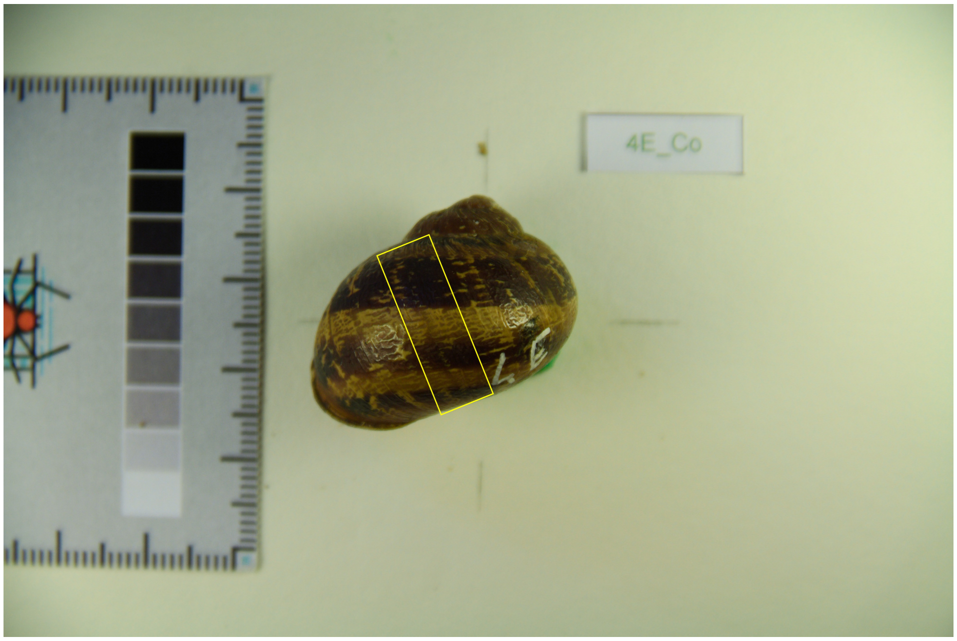


**Figure S2-1.** Example photograph used in reflectance analysis, showing both the snail shell in dorsal view and the grey standard card. The yellow rectangle highlights the area of the shell typically used in measurements.

We used the *errors* R package (Ucar, Pebesma, and Azcorra 2018) to correctly propagate the uncertainty from R, G, B channel predictions to their average via the Taylor series method, and accounted for it in our models (see **Statistical analyses** in the main text). Indeed, while the calibration curves had very good overall performance, we found that prediction uncertainty/residual standard deviation could not be neglected when these curves were applied to specifically *Cornu aspersum* shells. This is because, while prediction uncertainty was small compared to the entire range of the grey standard, *C. aspersum* shells are all relatively dark, i.e. with consistently small reflectance values. Indeed, estimated prediction uncertainty was on average 5.94% of estimated predicted value (range: 2.67% - 22.27%).

### S3 - Overview of models

General notes:

- All parameters interpretable as "fixed-effects" coefficients are denoted as $\beta$ (with intercepts as $\beta_0$), random intercepts as $\alpha$ or $\gamma$, while all parameters interpretable as standard deviations or standard errors are denoted as $\sigma$.
- For notation simplicity, variables names are reset between models, and the way missing values and observation error are dealt with is not always detailed (see McElreath 2020 for explicit information).
- All response variables in Gaussian models (so **Model 1** below), as well as all continuous predictors in all models, are assumed to be centred and scaled to unit 1 (observed) SD.

### Model 1 - Effect of urbanization on shell phenotypic traits

We can model the shell size (greater **D**iameter) $D_{i,j}$ of snail $j$ from site $i$ and their shell reflectance (**C**olour) $C_{i,j}$ as correlated variables. As shell size measurement error is extremely low (see main text **Methods**), we ignore it for modelling purposes. However, this is not the case for reflectance (see **Supplementary Material S2** above and file `00a-colour_calibration` in archived code for details); we therefore explicitly model the true (unknown) reflectance $C_{i,j}$ from the observed measurement $C_{i,j[obs]}$ and its associated measurement error $\sigma_{i,j[obs,C]}$ (McElreath 2020):

$$C_{i,j[obs]} \sim \text{Normal}(C_{i,j}, \sigma_{i,j[obs,C]}),$$

$$\begin{bmatrix} C_{i,j} \\ D_{i,j} \end{bmatrix} \sim \text{MVNormal}\left(\begin{bmatrix} \mu_{i[C]} \\ \mu_{i[D]} \end{bmatrix}, \Omega_{[residual]}\right),$$

$$\Omega_{[residual]} = \begin{bmatrix} \sigma_{[C]} & 0 \\ 0 & \sigma_{[D]} \end{bmatrix} R_{[residual]} \begin{bmatrix} \sigma_{[C]} & 0 \\ 0 & \sigma_{[D]} \end{bmatrix},$$

where $R_{[residual]}$ is the *residual* (here snail-level) correlation between the two response variables.

Expected values depend on predictors (here only urbanization $x_{1,i}$) and site identity as follows:

$$\mu_{i[D]} = \beta_{0[D]} + \beta_{1[D]} \times x_{1,i} + \alpha_{i[D]},$$

$$\mu_{i[C]} = \beta_{0[C]} + \beta_{1[C]} \times x_{1,i} + \alpha_{i[C]},$$

with the site-level random intercepts $\alpha_i$ for each response linked together through a common variance-covariance matrix $\Omega_\alpha$:

$$\begin{bmatrix} \alpha_{i[D]} \\ \alpha_{i[C]} \end{bmatrix} \sim \text{MVNormal}\left(\begin{bmatrix} 0 \\ 0 \end{bmatrix}, \Omega_\alpha\right),$$

$$\Omega_\alpha = \begin{bmatrix} \sigma_{\alpha[D]} & 0 \\ 0 & \sigma_{\alpha[C]} \end{bmatrix} R_\alpha \begin{bmatrix} \sigma_{\alpha[D]} & 0 \\ 0 & \sigma_{\alpha[C]} \end{bmatrix}.$$

We used $\text{Normal}(0,1)$ priors for the intercepts and fixed effects $\beta$, $\text{Half}-\text{Normal}(0,1)$ priors for all $\sigma$s, and $\text{LKJ}(2)$ priors for the correlation matrices $R$.

### Model 2 - Effect of urbanization and shell phenotype on parasites

We can model **S**porocysts presence/absence $S_{i,j}$, **M**etacercariae presence/absence $M_{i,j}$, *Nemhelix **b**akeri* presence/absence $B_{i,j}$, and the presence/absence of the **O**ther nematodes $O_{i,j}$ as follows:

$$\begin{aligned}
S_{i,j} &\sim \text{Bernoulli}(p_{i,j[S]}),\\
M_{i,j} &\sim \text{Bernoulli}(p_{i,j[M]}),\\
B_{i,j} &\sim \text{Bernoulli}(p_{i,j[B]}),\\
O_{i,j} &\sim \text{Bernoulli}(p_{i,j[O]}),\\
\text{logit}(p_{i,j[S]}) &= \beta_{0[S]} + \sum_{n=1}^{N}(\beta_{n[S]} \times x_{n,i,j}) + \alpha_{i[S]},\\
\text{logit}(p_{i,j[M]}) &= \beta_{0[M]} + \sum_{n=1}^{N}(\beta_{n[M]} \times x_{n,i,j}) + \alpha_{i[M]},\\
\text{logit}(p_{i,j[B]}) &= \beta_{0[B]} + \sum_{n=1}^{N}(\beta_{n[B]} \times x_{n,i,j}) + \alpha_{i[B]},\\
\text{logit}(p_{i,j[O]}) &= \beta_{0[O]} + \sum_{n=1}^{N}(\beta_{n[O]} \times x_{n,i,j}) + \alpha_{i[O]},
\end{aligned}$$

where $x_{n,i,j}$ are the snail values for each of the $N$ predictor variables, here urbanization, shell size, shell reflectance (for a view of how uncertainty in shell reflectance is accounted for when on the predictor side, see McElreath 2020). As in the previous model, random intercepts $\alpha$ are linked through a shared variance-covariance matrix:

$$\begin{bmatrix} \alpha_{i[S]} \\ \alpha_{i[M]} \\ \alpha_{i[B]} \\ \alpha_{i[O]} \end{bmatrix} \sim \text{MVNormal}\left( \begin{bmatrix} 0 \\ 0 \\ 0 \\ 0 \end{bmatrix}, \Omega_\alpha \right),$$

$$\Omega_\alpha = \begin{bmatrix} \sigma_{\alpha[S]} & 0 & 0 & 0 \\ 0 & \sigma_{\alpha[M]} & 0 & 0 \\ 0 & 0 & \sigma_{\alpha[B]} & 0 \\ 0 & 0 & 0 & \sigma_{\alpha[O]} \end{bmatrix} R_\alpha \begin{bmatrix} \sigma_{\alpha[S]} & 0 & 0 & 0 \\ 0 & \sigma_{\alpha[M]} & 0 & 0 \\ 0 & 0 & \sigma_{\alpha[B]} & 0 \\ 0 & 0 & 0 & \sigma_{\alpha[O]} \end{bmatrix}.$$

We used $\text{Normal}(0, 1.5)$ priors for the intercepts $\beta_0$ (which are interpretable as the logit of proportions, McElreath 2020), $\text{Normal}(0, 1)$ priors for the other fixed effects $\beta$, $\text{Half} - \text{Normal}(0, 1)$ priors for random effect SDs $\sigma$, and an $\text{LKJ}(2)$ prior for the correlation matrix $R$.

### Model 3 - Effect of urbanization, shell phenotype and parasites on behaviour

Finally, we can model the proportion of **F**ood consumed $F_{i,j,k}$ (site $i$, snail $j$, trial $k$)[1] and movement **A**ctivity $A_{i,j,k}$ as follows:

$$F_{i,j,k} \sim \text{Beta}(p_{i,j[F]}, \phi_{[F]}),$$
$$A_{i,j,k} \sim \text{Weibull}(\mu_{i,j[A]}, \theta_{[A]}),$$

where $\phi$ is a precision parameter, and $\theta$ a shape parameter (see https://cran.r-project.org/web/packages/brms/vignettes/brms_families.html [accessed 2024-10-16]).

Formula for the means for each response can be written as:

$$\text{logit}(p_{i,j[F]}) = \beta_{0[F]} + \sum_{n=1}^{N} (\beta_{n[F]} \times x_{n,i,j}) + \alpha_{i[F]} + \gamma_{j[F]},$$

$$\log(\mu_{i,j[A]}) = \beta_{0[A]} + \sum_{n=1}^{N} (\beta_{n[A]} \times x_{n,i,j}) + \alpha_{i[A]} + \gamma_{j[A]},$$

with here random effects for both the site ($\alpha_i$) and snail ($\gamma_j$) levels due to repeated measurements. Both site- and snail-level random effects have their own variance-covariance matrix:

$$\begin{bmatrix} \alpha_{i[F]} \\ \alpha_{i[A]} \end{bmatrix} \sim \text{MVNormal}\left( \begin{bmatrix} 0 \\ 0 \end{bmatrix}, \Omega_\alpha \right),$$

$$\begin{bmatrix} \gamma_{i[F]} \\ \gamma_{i[A]} \end{bmatrix} \sim \text{MVNormal}\left( \begin{bmatrix} 0 \\ 0 \end{bmatrix}, \Omega_\gamma \right),$$

$$\Omega_\alpha = \begin{bmatrix} \sigma_{\alpha[F]} & 0 \\ 0 & \sigma_{\alpha[A]} \end{bmatrix} R_\alpha \begin{bmatrix} \sigma_{\alpha[F]} & 0 \\ 0 & \sigma_{\alpha[A]} \end{bmatrix},$$

$$\Omega_\gamma = \begin{bmatrix} \sigma_{\gamma[F]} & 0 \\ 0 & \sigma_{\gamma[A]} \end{bmatrix} R_\gamma \begin{bmatrix} \sigma_{\gamma[F]} & 0 \\ 0 & \sigma_{\gamma[A]} \end{bmatrix}.$$

[1] proportions transformed to avoid issues with the few zeroes present, following Douma and Weedon (2019) (see main text).

We set a Normal$(0, 1.5)$ prior for $\beta_{0[F]}$ (interpretable as the logit of a proportion, see **Model 2**) and Normal$(0, 1)$ priors for all other $\beta$s. As with previous models, we used Half $-$ Normal$(0, 1)$ priors for random effect SDs $\sigma$ and an LKJ(2) prior for the correlation matrices $R$. We set a Exponential(1) prior on $\theta$. For the Beta precision parameter $\phi$, we follow McElreath (2020) in setting a Exponential(1) prior on $(\phi - 2)$ rather than $\phi$ itself.

### S4 - Model posteriors

In the main text, only parameter summaries in the form "mean [95% quantile interval]" are given, for simplicity. We provide below the corresponding plots showing the full posterior distributions.

#### Model 1 - Effect of urbanization on shell phenotypic traits

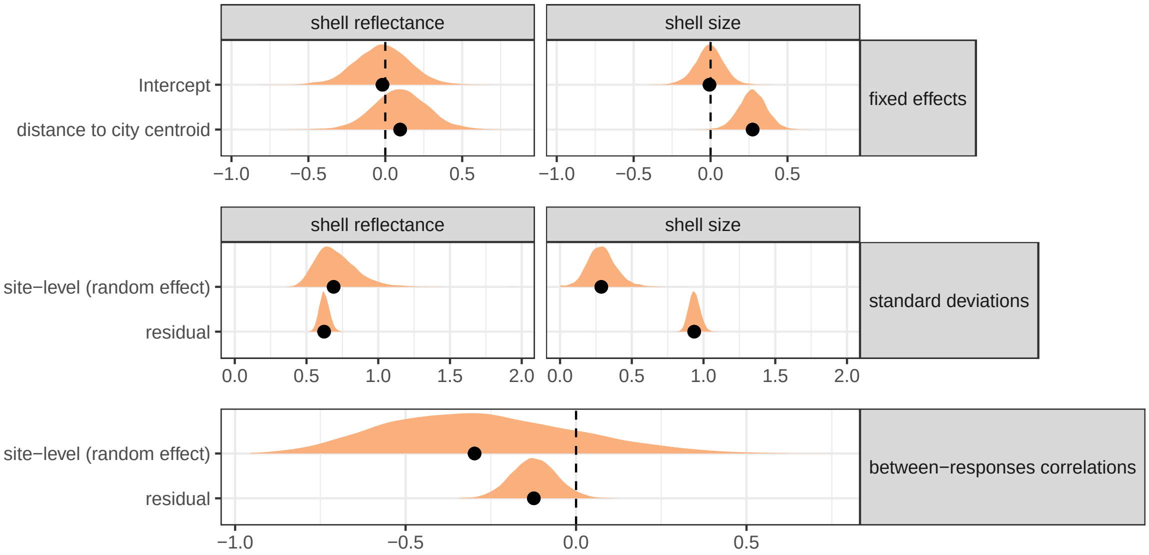


**Figure S4-1.** Posterior densities for the "shell phenotype" model.

### Model 2 - Effect of urbanization and shell phenotype on parasites

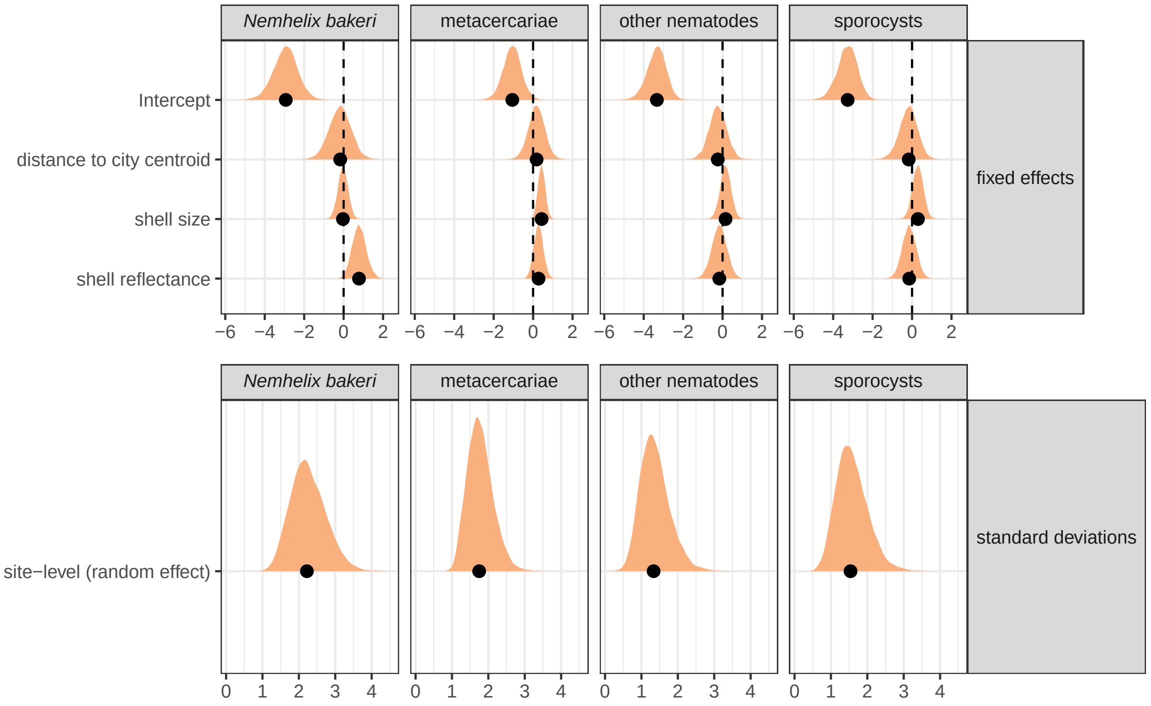


**Figure S4-2.** Posterior densities for the fixed effects and random effect SDs of the parasite prevalence model.

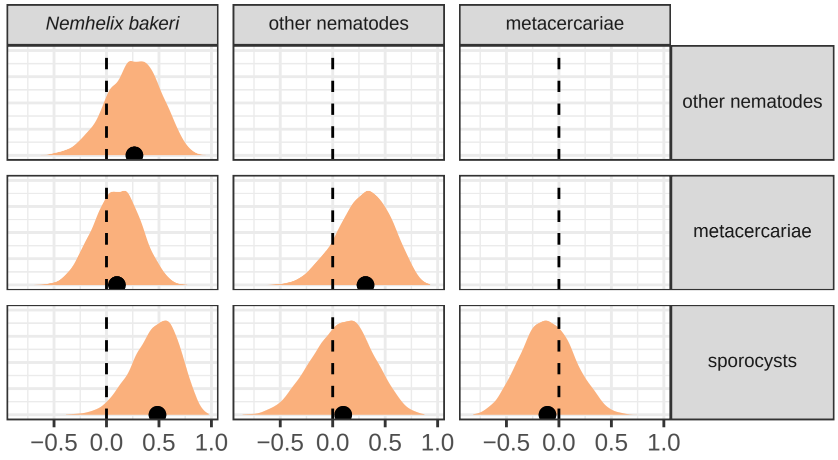


**Figure S4-3.** Posterior densities for the site-level correlations between responses in the parasite prevalence model.

### Model 3 - Effect of urbanization, shell phenotype and parasites on behaviour

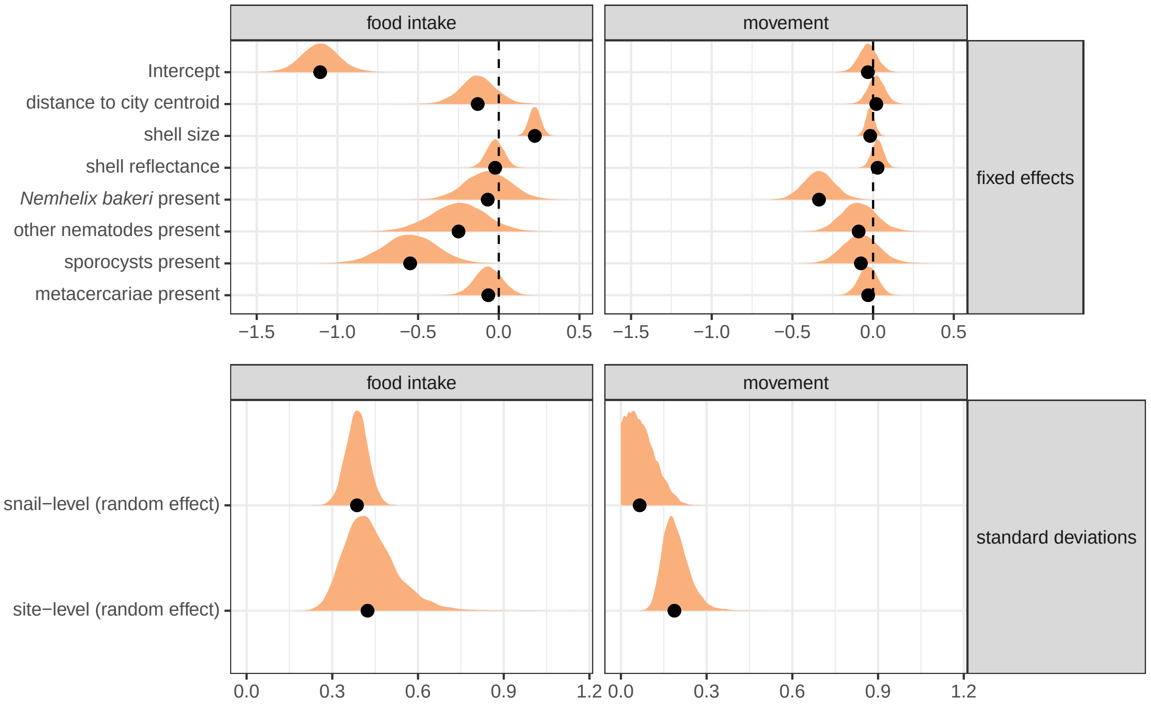


**Figure S4-4.** Posterior densities for the fixed effects and random effect SDs of the behaviour model.

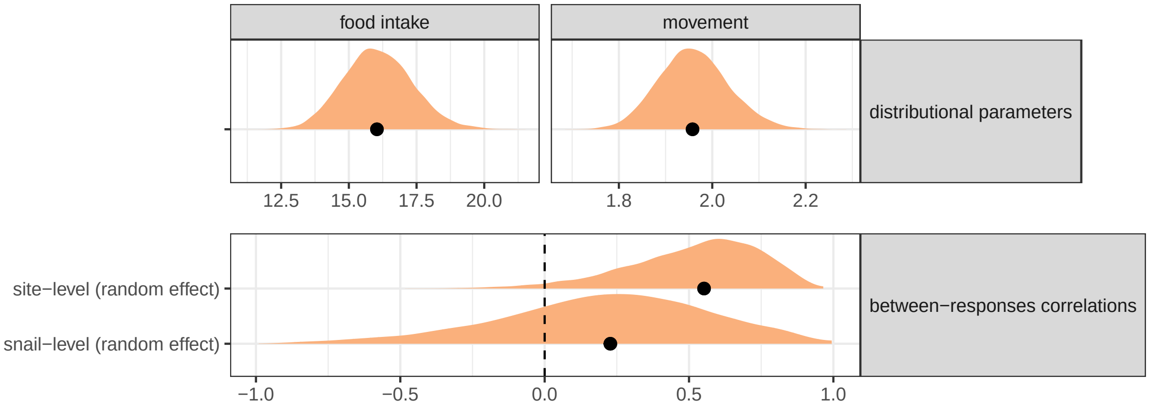


**Figure S4-5.** Posterior densities for the distributional parameters and correlations between responses of the parasite prevalence model. Distributional parameters are a precision parameter $\phi$ for the food intake submodel and a shape parameter $\theta$ for the movement model (see **Supplementary Material S3** for details).

### S5 - Overview of parasite prevalences

**Table S5-1.** Observed prevalences of each parasite type in sampled *Cornu aspersum* snails, both per site and overall. Sites are ordered by increasing distance from the city centroid ("Distance (m)" column).

| | | | | Prevalence of: | | | |
|---|---|---|---|---|---|---|---|
| Site ID | Distance (m) | Longitude | Latitude | Sporocysts | Metacercariae | *N. bakeri* | Other nematodes |
| 20 | 1519 | 1°40'55"W | 48°6'24"N | 0.00 (0/16) | 0.69 (11/16) | 0.19 (3/16) | 0.12 (2/16) |
| 10 | 1714 | 1°41'10"W | 48°6'54"N | 0.00 (0/20) | 0.00 (0/20) | 0.00 (0/20) | 0.00 (0/20) |
| 11 | 1728 | 1°40'37"W | 48°7'31"N | 0.15 (3/20) | 0.00 (0/20) | 0.00 (0/20) | 0.00 (0/20) |
| 2 | 1951 | 1°41'21"W | 48°6'32"N | 0.00 (0/20) | 0.10 (2/20) | 0.00 (0/20) | 0.05 (1/20) |
| 18 | 2031 | 1°41'8"W | 48°7'24"N | 0.08 (1/12) | 0.17 (2/12) | 0.33 (4/12) | 0.08 (1/12) |
| 9 | 2358 | 1°41'29"W | 48°7'21"N | 0.15 (3/20) | 0.60 (12/20) | 0.05 (1/20) | 0.00 (0/20) |
| 3 | 2385 | 1°41'43"W | 48°6'38"N | 0.00 (0/18) | 0.28 (5/18) | 0.00 (0/18) | 0.00 (0/18) |
| 4 | 2872 | 1°42'7"W | 48°6'48"N | 0.05 (1/20) | 0.65 (13/20) | 0.05 (1/20) | 0.40 (8/20) |
| 19 | 2957 | 1°41'56"W | 48°7'28"N | 0.10 (1/10) | 0.30 (3/10) | 0.70 (7/10) | 0.10 (1/10) |
| 13 | 3391 | 1°41'50"W | 48°7'58"N | 0.00 (0/20) | 0.45 (9/20) | 0.00 (0/20) | 0.00 (0/20) |
| 12 | 3591 | 1°42'26"W | 48°7'35"N | 0.00 (0/19) | 0.16 (3/19) | 0.00 (0/19) | 0.05 (1/19) |
| 7 | 3660 | 1°42'13"W | 48°7'53"N | 0.00 (0/16) | 0.06 (1/16) | 0.00 (0/16) | 0.00 (0/16) |
| 6 | 4182 | 1°42'39"W | 48°7'58"N | 0.50 (10/20) | 0.10 (2/20) | 0.45 (9/20) | 0.05 (1/20) |
| 5 | 4316 | 1°42'45"W | 48°7'59"N | 0.00 (0/20) | 0.90 (18/20) | 0.00 (0/20) | 0.00 (0/20) |
| 16 | 5006 | 1°43'5"W | 48°8'20"N | 0.06 (1/17) | 0.18 (3/17) | 0.00 (0/17) | 0.00 (0/17) |
| 14 | 5250 | 1°43'25"W | 48°8'13"N | 0.00 (0/18) | 0.06 (1/18) | 0.17 (3/18) | 0.00 (0/18) |
| 17 | 5918 | 1°43'51"W | 48°8'26"N | 0.00 (0/17) | 0.65 (11/17) | 0.00 (0/17) | 0.06 (1/17) |
| **Overall** | | | | | | | |
| — | — | — | — | 0.07 (20/303) | 0.32 (96/303) | 0.09 (28/303) | 0.05 (16/303) |

### S6 - Investigating the site-level correlation between reflectance and *N. bakeri* prevalence

In the main text, we ran analyses where individual shell traits were used as predictors for individual parasite prevalence. We found among other things that shell reflectance was linked to *Nemhelix bakeri* prevalence (main text **Fig. 3**). At the same time, we found that for both reflectance and *N. bakeri* prevalence, a substantial proportion of the total variance was at the population/site level.

This raises a question that our original models were not designed to address. Is the link between *Nemhelix bakeri* prevalence and reflectance due to individual-level correlations (i.e. even within a given population, lighter snails are more likely to be infected)? Or is it due to site-level correlations (sites with higher prevalence are also sites with on average lighter shells, possibly due to correlational selection, but there is no true link between the two at the individual level)?

To explore this, we ran a new bivariate model where both reflectance and *N. bakeri* prevalence were responses, correlated with each other through site-level random effects. Submodel families, and the way uncertainty in reflectance was dealt with, were the same as in the main models; no fixed effects were included. This new model allowed us to estimate the site-level correlation between the two responses (note that the residual individual-level correlation cannot be estimated from such a model due to prevalence being a binary response), and can be written as follows (see **Supplementary Material S3** for notation details):

$$
\begin{aligned}
B_{i,j} &\sim \text{Bernoulli}(p_{i[B]}),\\
C_{i,j[obs]} &\sim \text{Normal}(C_{i,j}, \sigma_{i,j[obs,C]}),\\
C_{i,j} &\sim \text{Normal}(\mu_{i[C]}, \sigma_{[C]}),\\
\text{logit}(p_{i[B]}) &= \beta_{0[B]} + \alpha_{i[B]},\\
\mu_{i[C]} &= \beta_{0[C]} + \alpha_{i[C]},\\
\begin{bmatrix}\alpha_{i[B]}\\ \alpha_{i[C]}\end{bmatrix} &\sim \text{MVNormal}\left(\begin{bmatrix}0\\0\end{bmatrix}, \Omega\right),\\
\Omega &= \begin{bmatrix}\sigma_{\alpha[B]} & 0\\ 0 & \sigma_{\alpha[C]}\end{bmatrix} R \begin{bmatrix}\sigma_{\alpha[B]} & 0\\ 0 & \sigma_{\alpha[C]}\end{bmatrix},
\end{aligned}
$$

with priors also as in **Supplementary Material S3**, models 1 & 2.

Running this model gives an estimate for the among-sites correlation of -0.30 [-0.70, 0.18], which is not different from zero (**Fig. S6-1**). While this does not necessarily mean that the link we uncovered in the main text is primarily due to *within-population* correlations (especially given the low number of sites with non-zero prevalence, **Supplementary Material S5**), it does suggest this may be the case.

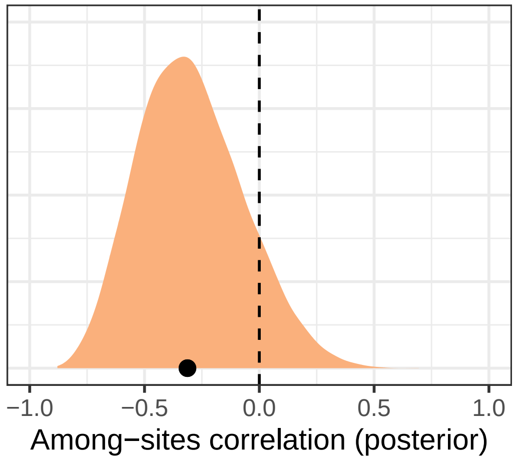


**Figure S6-1.** Posterior distribution of the site-level correlation between *N. bakeri* prevalence and shell reflectance.

McElreath, Richard. 2020. *Statistical Rethinking: A Bayesian Course with Examples in R and Stan*. 2nd edition. Boca Raton, USA: Chapman and Hall/CRC.

Schindelin, Johannes, Ignacio Arganda-Carreras, Erwin Frise, Verena Kaynig, Mark Longair, Tobias Pietzsch, Stephan Preibisch, et al. 2012. "Fiji: An Open-Source Platform for Biological-Image Analysis." *Nature Methods* 9 (7): 676–82. https://doi.org/10.1038/nmeth.2019.

Ucar, Iñaki, Edzer Pebesma, and Arturo Azcorra. 2018. "Measurement Errors in R." *The R Journal* 10 (2): 549–57. https://doi.org/10.32614/RJ-2018-075.

Williamson, Phillip. 1979. "Age Determination of Juvenile and Adult *Cepaea*." *Journal of Molluscan Studies* 45: 52–60.
